# The Poisson-lognormal model as a versatile framework for the joint analysis of species abundances

**DOI:** 10.1101/2020.10.07.329383

**Authors:** Julien Chiquet, Mahendra Mariadassou, Stéphane Robin

**Affiliations:** MIA-Paris, Université Paris-Saclay – AgroParisTech – INRAE, Paris, France; MaIAGE, Université Paris-Saclay – INRAE, Jouy-en-Josas, France; CESCO, UMR 7204, MNHN - CNRS - UPMC, Paris, France

**Keywords:** abundance data, joint species distribution model, latent variable model, multivariate analysis, variational inference, R package

## Abstract

Joint Species Abundance Models (JSDM) provide a general multivariate framework to study the joint abundances of all species from a community. JSDM account for both structuring factors (environmental characteristics or gradients, such as habitat type or nutrient availability) and potential interactions between the species (competition, mutualism, parasitism, etc.), which is instrumental in disentangling meaningful ecological interactions from mere statistical associations.

Modeling the dependency between the species is challenging because of the count-valued nature of abundance data and most JSDM rely on Gaussian latent layer to encode the dependencies between species in a covariance matrix. The multivariate Poisson-lognormal (PLN) model is one such model, which can be viewed as a multivariate mixed Poisson regression model. The inference of such models raises both statistical and computational issues, many of which were solved in recent contributions using variational techniques and convex optimization.

The PLN model turns out to be a versatile framework, within which a variety of analyses can be performed, including multivariate sample comparison, clustering of sites or samples, dimension reduction (ordination) for visualization purposes, or inference of interaction networks. This paper presents the general PLN framework and illustrates its use on a series a typical experimental datasets. All the models and methods are implemented in the `R` package `PLNmodels`, available from cran.r-project.org.

## 1 Introduction

### 1.1 Joint species distribution models

Joint Species Distribution Models (JSDM) have received a lot of attention in the last decade as they provide a general multivariate framework to study the joint abundances of all species from a community, as opposed to species distribution models (SDM: Elith & Leathwick, 2009) where species are considered as disconnected entities. At their best, JSDM account for both structuring factors (e.g. environmental gradients, nutrients availaibility, etc) and potential interactions between the species (competition, mutualism, parasitism, etc.). Broadly speaking, such models include both abiotic and biotic effects to describe the fluctuations of species abundances across space and time. Considering both effects at once is instrumental in disentangling meaningful ecological interactions from mere statistical associations induced by environmental drivers and/or habitat preferences. JSDMs have been proposed to deal with presence/absence data (see for example Ovaskainen *et al*., 2010; Harris, 2015), for abundance data (Popovic *et al*., 2019), or both (Popovic *et al*., 2018). We focus here on abundance data, and more specifically on data which consists of a count associated with each species in each site, date or condition.

Modeling the dependency between the species is challenging because of the count-valued nature of abundance data. Contrarily to continuous multivariate distributions, there exists no generic multivariate distribution for count data (Inouye *et al*., 2017). As a consequence, many JSDM rely on the same hierarchical backbone: dependencies are first modeled in a latent layer through the covariance matrix of a multivariate, most often Gaussian, vector and counts are then sampled independently conditionally to this latent (Gaussian) vector of expected (transformed-)abundances (see Warton *et al*., 2015, for a general presentation). Dependencies between counts are fully captured by the covariance matrix of the latent vector, whereas environmental effects are accounted for in the vector mean value. This distinction is convenient from a modeling point of view, as it typically separates a regression part (taking the point of view of multivariate generalized linear model) that accounts for abiotic effects, and a random part that accounts for dependency between species (biotic effects).

This paper introduces the Poisson-lognormal (PLN) model – first proposed by Aitchison & Ho (1989) – as a JSDM. Broadly speaking, the PLN model can be viewed as a multivariate mixed generalized linear model with Poisson distribution. Because of its simple form, the PLN model turns out to be versatile in the sense that it provides a convenient framework to carry out a series of usual (multivariate) statistical analyses. This includes multivariate regression in its simplest form, but also multivariate sample comparison via linear discriminant analysis (LDA), model-based clustering using mixture models, dimension reduction via principal component analysis (PCA: Chiquet *et al*., 2018), and network inference (Chiquet *et al*., 2019). All these analyses are implemented in the `R` package `PLNmodels`, available from cran.r-project.org. Because it involves a latent layer, the inference of this model raises a series of computational issues, which can be circumvented via a variational approximation (Blei *et al*., 2017).

The rest of this section is devoted to the precise definition of the PLN model. Section 2 gives a brief introduction to the variational inference approach implemented in the `PLNmodels` package and how measures of uncertainty for the parameters can be derived from it. Section 3 provides a series of examples illustrating how the PLN model can be adapted to tackle some specific questions (sample comparison, clustering, dimension reduction and visualization, or network inference) using various extensions summarized in Figure 1. The last section provides additional information about the `PLNmodels` package and describes several research leads motivated by current needs in ecological modeling.

**Figure 1:**
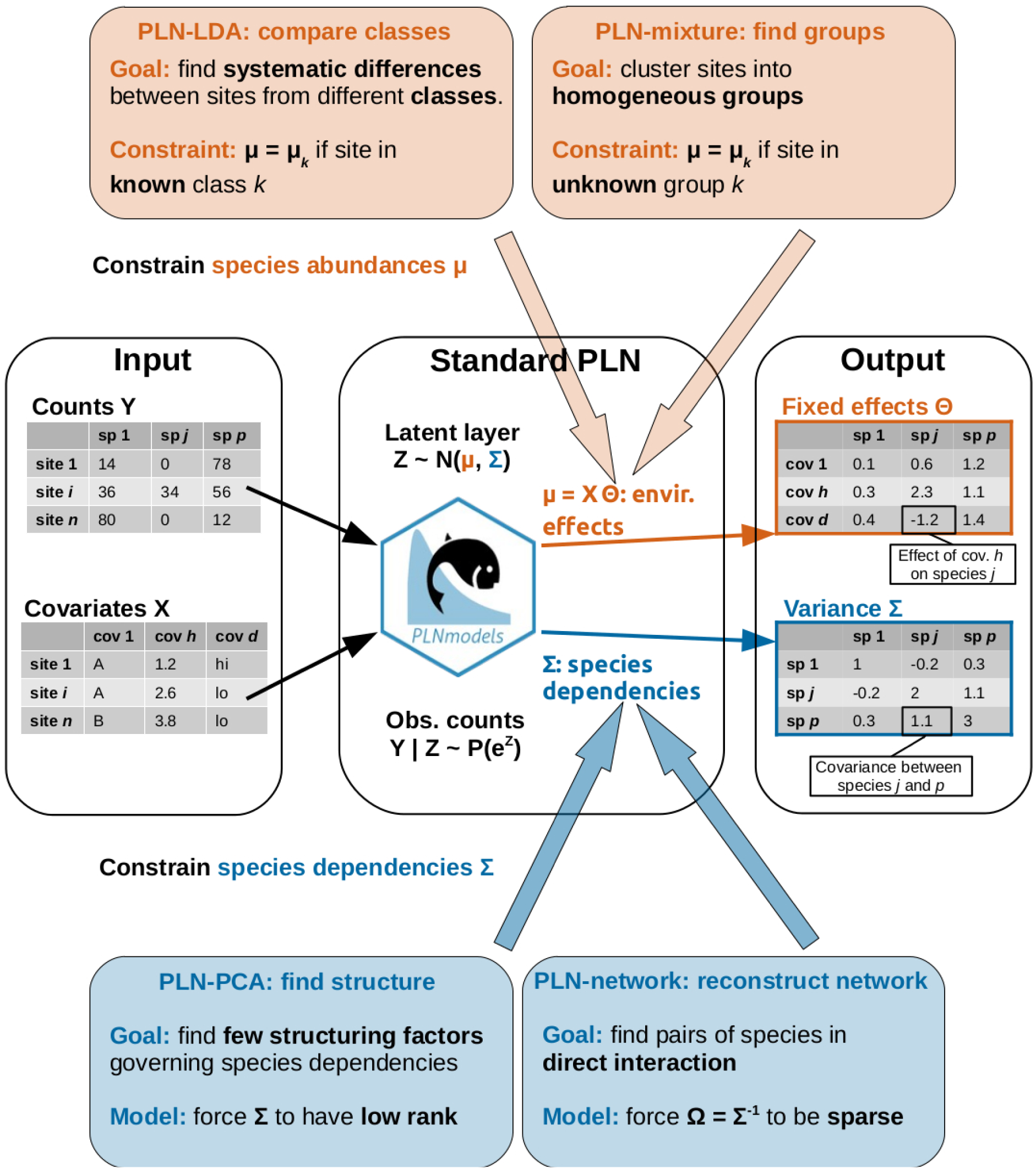
Graphical abstract of the article: the standard PLN model, its extension and the research questions addressed by each method.

### 1.2 The Poisson-lognormal model

The multivariate Poisson-lognormal model (Aitchison & Ho, 1989) is designed for the analysis of an abundance table, that is typically a *n* × *p* count matrix *Y*, where *Y*_*ij*_ is the number of individuals from species *j* observed in site *i*, n being the number of sites and p the number of species. Note that ’site’ may actually refer to a sample or an experiment, and a ’species’ to an Operational Taxonomic Unit (OTU) or an Amplicon Sequence Variant (ASV), both of which are proxies for species frequently used in metabarcoding surveys. Similarly, the ’number of individuals’ may correspond to a number of reads in a metabarcoding experiments.

The PLN models relates the p-dimensional abundance vector *Y*_*i*_ collected in site *i* with a *p*-dimensional Gaussien latent vector *Z*_*i*_ as follows:

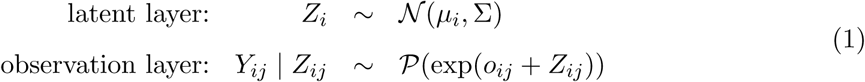

where the *Z*_*i*_ are supposed to be independent (independence across sites) and the abundances *Y*_*ij*_ are all conditionally independent given the latent variables *Z*_*ij*_. The parameter *µ*_*i*_ = [*µ*_*ij*_]_1≤*j*≤*p*_ ∈ ℝ^*p*^ corresponds to the fixed effects, and is related to the expected log-abundances, whereas the latent covariance matrix Σ = [*σ*_*jk*_]_1≤*j,k*≤*p*_ describes the underlying structure of dependence between the p species. In this simple form, the PLN model therefore assumes that the dependency structure among species is the same across all sites. All extensions presented in Section 3 will be about alternative modeling of the latent layer in Eq. 1, corresponding to different assumptions exposed in Figure 1.

When environmental covariates are available, the fixed effect *µ*_*i*_j in the latent layer may be decomposed as 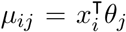 where *x*_*i*_ ∈ ℝ^*d*^ is a vector of covariates for sample i (e.g. environmental conditions, location, etc.) and *θ*_*j*_ ∈ ℝ^*d*^ is a vector of regression coefficients associated to these *d* covariates for species *j*. The fixed quantity *o*_*ij*_ is the offset for species *j* in sample *i*. In the PLN framework, offsets are used to take into account expected differences in observed counts due to imbalanced sampling efforts, such as known heterogeneities in terms, sequencing depths in metabarcoding surveys, collection protocols, species detectability, etc. For examples, if we spend twice as much time looking for species *j* in site *i′* than in site *i* and thus expect its count to be twice higher (all other things being equal), we set *o*_*i′j*_ = *o*_*ij*_ + log(2).

Likewise in generalized linear models, the parameters should be interpreted according to the properties of the multivariate Poisson-lognormal distribution. Some remarks can be made about the first and second order moments, which are given by Aitchison & Ho (1989):

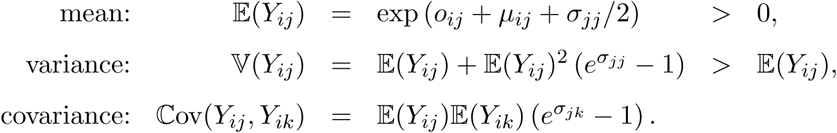

#### Expected count

Due to the ’log’ link function, the expected abundance 𝔼(*Y*_*ij*_) of species *j* in site *i* is not simply exp(*o*_*ij*_ + *µ*_*ij*_) as the variance parameter *σ*_*jj*_ is also involved.

#### Over-dispersion

Because of the presence of a latent (random) layer, the PLN model displays a larger variance than the Poisson model for which 𝕍(*Y*_*ij*_) = 𝔼(*Y*_*ij*_).

#### Faithful correlation

Because 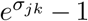 has the same sign as *σ*_*jk*_, the covariance (resp. correlation) between the respective abundances *Y*_*ij*_ and *Y*_*ik*_ of species *j* and *k* has the same sign as the covariance (resp. correlation) between the corresponding latent components *Z*_*ij*_ and *Z*_*ik*_.

The last property is especially desirable as it means that the correlation structure of the latent vector *Z*_*i*_ preserves the one of the observed abundances *Y*_*i*_. As a consequence, the independence of *Z*_*ij*_ and *Z*_*ik*_ (*σ*_*jk*_ = 0) induces an absence of correlation between *Y*_*ij*_ and 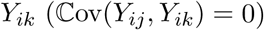.

## 2 Parameter estimation

### 2.1 Inference algorithm

#### 2.1.1 Variational inference

Because the latent layer *Z* is not observed, the PLN model is an incomplete data model in the sense of Dempster *et al*. (1977), who introduced the celebrated EM algorithm to perform maximum likelihood for such models. This is an iterative two-steps algorithm. Intuitively, the E step retrieves, from the observed counts *Y*_*i*_, all the information about the latent vectors *Z*_*i*_’s that is needed, during the M step, to estimate the parameters Θ and Σ. More formally, the E step requires the evaluation of the conditional distribution of the latent vectors given the observed counts, that is: *p*(*Z* | *Y*). Unfortunately, this distribution is intractable for the PLN model, so we resort to a variational approximation (see e.g. Jaakkola, 2001) of this conditional distribution. This results in what is called a variational EM (VEM) algorithm, which alternates the following two steps until convergence:

##### VE step

Given the current estimates 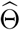 and 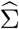 of the parameters, for each site *i*, find the normal distribution 𝒩 (*m*_*i*_, *S*_*i*_) that best fits the (unknown) conditional distribution *p*(*Z*_*i*_ | *Y*_*i*_) in terms of Küllback-Leibler (KL) divergence;

##### M step

Given the approximate conditional distribution of the latent *Z*_*i*_’s, update the parameter estimates 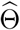 and 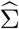.

The approximation precisely lies in the fact that the true conditional distribution *p*(*Z*_*i*_ | *Y*_*i*_) is not Gaussian. Hence, the approximate distribution *q*(*Z*) is a product over the sites of Gaussian distributions 𝒩 (*Z*_*i*_; *m*_*i*_, *S*_*i*_). The approximate mean *m*_*i*_ and the (diagonal) covariance matrix *S*_*i*_ are called the *variational parameters*; they can be merged into two *n* × *p* matrices, denoted *M* and *S*, respectively. Using such an approximation amounts to maximizing a lower bound of the log-likelihood of the data log *p*(*Y*; Θ, Σ) (Blei *et al*., 2017):

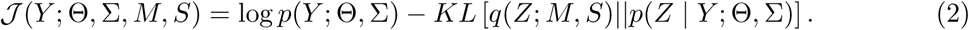

The objective function of the VEM therefore depends on the data *Y* and has to be optimized with respect to both the model parameters (Θ, Σ) and the variational parameters (*M, S*).

#### 2.1.2 A ftrst example

As a first illustration of the use of the PLN model for analyzing abundance data, we consider the dataset introduced by Fossheim *et al*. (2006) (and re-analyzed by Greenacre, 2013; Greenacre & Primicerio, 2014), which consists of the abundances of *p* = 30 fish species measured in n = 89 sites of the Barents sea between April and May 1997. The species under study are sensitive to environmental drivers (temperature, water depth,...) but are also related by trophic interactions. A first aim is to study the relative contribution of each type of interaction (abiotic vs biotic) to the structure of the community.

Captures were carried out with the same protocol for all species and all sites, so no offset term is required here. For each site i, four covariates were recorded: the latitude, the longitude, the depth and the temperature, which constitute the vector xi. We also add an intercept to these covariates to capture differences in the base abundances of our 30 species. The covariates can be gathered into a *n* × (*d* = 5) matrix *X*.

Fitting the PLN model results in a matrix of regression coefficients 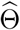 and a latent covariance matrix 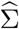. Figure 2 shows these estimates. The most contrasted regression coefficient turns out to be the intercept, revealing a great variability between the mean abundances of the species. The latent correlation structure encoded in 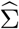 reveals some subgroups of species: the covariation between these species are not caused by the effects of the recorded covariates.

**Figure 2:**
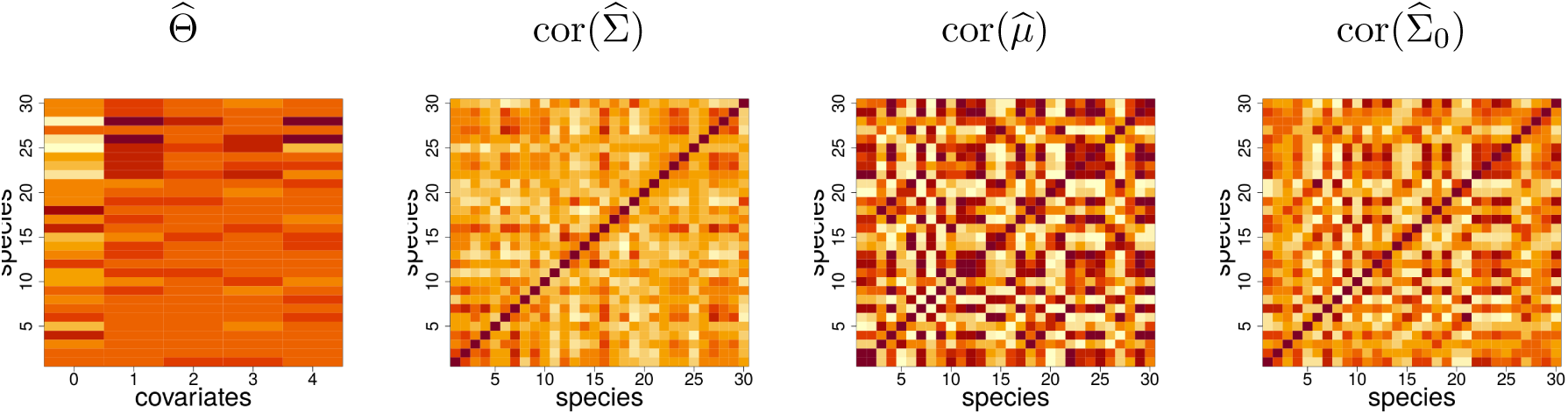
Fit of the PLN model to the Barents fish data set. 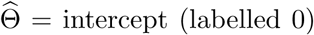 and regression coefficients for the four covariates and the 30 species; 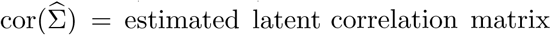; 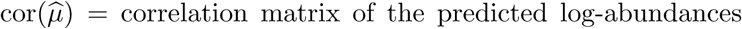; 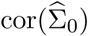 = estimated correlation matrix for the null model, which includes only an intercept term.

Based on this, a predicted log-abundance can be computed for each species at each site as 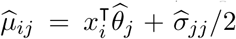. Interestingly the correlation structure between these predicted log-abundances is very constrasted compared to 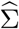. This shows that a substantial part of the covariation between species abundances is driven by the covariates. To illustrate this point, we fitted a model with only an intercept term, yielding the null covariance matrix 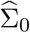, which captures the covariation between species due to both biotic and abiotic effects. We observe that 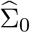 displays a structure quite similar to the correlation matrix of the prediction 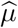, which shows the predominant contribution of environmental effects to empirical covariations.

### 2.2 Parameter uncertainty

Variational inference is useful to estimate parameters in the PLN model as it allows us to bypass the intractable likelihood. Unfortunately, this convenience comes at a cost as there are no *out of the box* theoretical guarantees on the quality of the estimates (unbiasedness, efficiency, asymptotic normality, etc.). We use large scale simulations to study the empirical properties of the variational estimates of the PLN model.

#### 2.2.1 Simulation settings

We simulated count data according to a PLN model with the following parameters: number of samples *n* ∈ {50, 100, 500, 1000, 10000}, number of species *p* ∈ {20, 200}, number of covariates *d* ∈ {2, 5, 10}, sampling effort *N* ∈ {low, medium, high} (calibrated to roughly correspond to total sums of counts per sample of 10^4^, 10^5^ and 10^6^), and noise level *σ*^2^ ∈ {0.2, 0.5, 1, 2}. These parameters cover values typically observed in real datasets and range from very hard (*n* = 50, *p* = 200, *d* = 10, *N* = low) to ridiculously easy (*n* = 10000, *p* = 20, *d* = 2, *N* = high).

For each of the 360 parameter combinations, hereafter referred to as *simulation setup*, we generated a variance matrix Σ as *σ*_*jk*_ = *σ*^2^*ρ*^|*j*−*k*|^, with *ρ* = 0.2, a design matrix *X* with 1s in the first column (intercept) and all other entries sampled from a standard Gaussian distribution (we also centered all columns but the first one to avoid interplay between the intercept and the covariates), a regression coefficient matrix Θ with all entries sampled from a centered Gaussian distribution with variance 1/*d*. Those choices ensures (*i*) a moderate correlation of the counts across species and (*ii*) the same order of magnitude for the fixed effects (*X*Θ) and the biological variability of the species (*σ*^2^).

For each simulation setup, we generated *R* = 100 count matrices (*Y*^(1)^,…, *Y*^*(R)*^), resulting in a total of 36 000 data sets. A PLN model was then fitted to each of those, resulting in *R* estimates 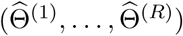 for each original matrix Θ.

#### 2.2.2 Bias

For each simulation setup and each coefficient *θ*_*jh*_ of that setup (were *j* refers to the species and *h* to the covariate, including the intercept), we computed the empirical bias as 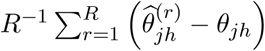 and the Root Mean Squared Error (RMSE) as 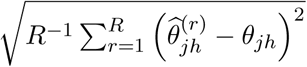. The distribution of those metric are represented in Fig. 3. Panel *a*) represents boxplots of the bias (one boxplot per setup, computed over all entries of Θ). It shows that the variational estimates are unbiased and, as expected, that the empirical bias decreases when the number of samples (*n*) increases and when the variability (*σ*^2^) decreases (note the different y-axis scales for different values of *σ*^2^). Panel *b*) likewise shows that the RMSE decreases with increasing *n* (as 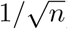) and decreasing values of *σ*^2^ (note again the different y-axis scales). By contrast, the sampling effort has no effect on the accuracy of the estimates. Considering that the typical scale for coefficient *θ*_*jh*_ is 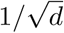, the RMSE is quite small (below 0.05) as soon as *n* ≥ 500 sites, for all values of *σ*^2^.

**Figure 3:**
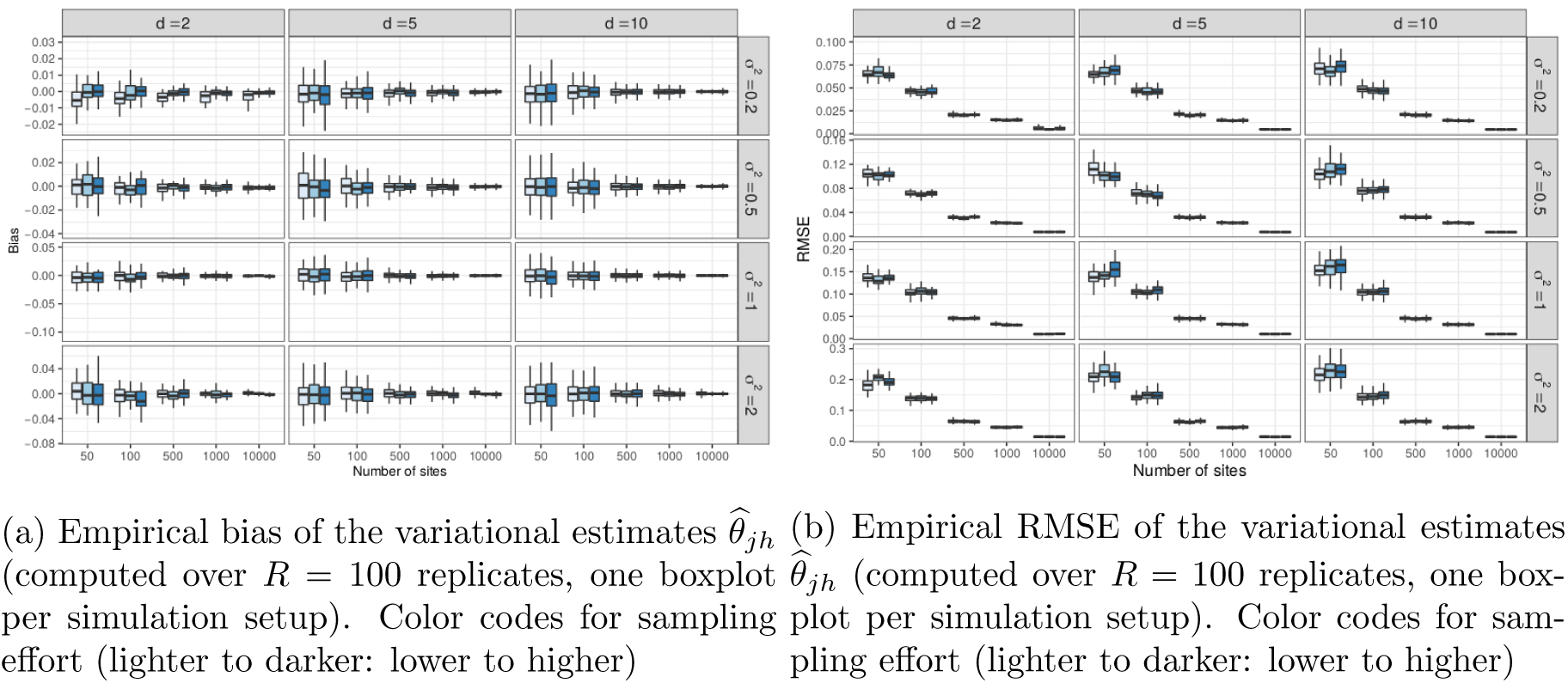
Result of the simulations: variational estimates are unbiased and have low RMSE starting at *n* = 500 (and even *n* = 100 for low levels of variability).

#### 2.2.3 Conftdence intervals

Few general theoretical results exist about the statistical properties of variational estimates, but naive approaches are known to provide too narrow confidence intervals (see, e.g., Wang & Titterington, 2005; Westling & McCormick, 2015). One naive approach consists in computing the Fisher information matrix (and deduce the confidence intervals) of the model parameters 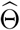 and 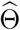 from the variational lower bound of Eq. (2), i.e. as if it were the true log-likelihood and not an approximation.

We used our simulation results to study the accuracy of confidence intervals computed this way. More specifically, we computed a 95% confidence interval 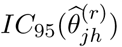 based on the pseudo-Fisher information matrix for each replicate r and computed their empirical coverage, that is the proportion of replicates for which the true parameter *θ*_*jh*_ lies within 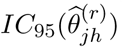. If the confidence intervals are well calibrated, this proportion should be around 0.95.

The results (not shown here) paint a striking picture as the coverage barely reaches 0.3 for low variability setups (*σ*^2^ = 0.2) and falls below 0.1 in more difficult setups, especially when the variability is large. They also show that higher sampling efforts do not improve the coverage. This counter intuitive finding confirms that the naive approximation does not provide reliable confidence intervals. Further developments are obviously needed to improve these shortcomings and some research leads are discussed in Section 4.

## 3 Illustrations

We now introduce a series of extensions of the basic PLN model defined in Eq. (1). As explained above, the PLN model deals with both abiotic effects through the mean vector *µ*_*i*_ and biotic effects by describing species interactions through the variance matrix Σ. As shown in Figure 1, the first two extensions (LDA and clustering) deal with the former and typically aim at analyzing abiotic (or environmental) effects. The last two extensions (PCA and network inference) are about the latter and provide insights about the dependency structure between the species.

Each method will be accompanied with a specific example. LDA will be used to compare the bacterial communities collected in different body sites of dairy cattle. Model-based clustering will be applied to the microbiota of leaves from several oaks and will prove to be able to recover, *in a blind way*, the tree of origin of each leaf. Dimension reduction using PCA will be applied to the same dataset for vizualization purposes and to exhibit which species contribute most to the community structure. The last example is an attempt to reconstruct the interaction network of fish species from the Barents Sea.

### 3.1 Sample comparison with linear discriminant analysis

A first variant of the PLN model is the analogue of the Linear Discriminant Analysis (LDA) to Poisson-lognormal models. It applies to labeled data, *i.e*. when sites belong to known classes (*e.g*. anatomical sites for host-associated microbiota, or geographical areas in ecogeography) and the objective is two-fold: identify class-based differences in species counts and predict the class of an unlabeled site based on its species counts. As valuable byproducts, classification accuracy assesses whether the classes are really different and species with a high contribution to the discriminant axes can serve as biomarkers.

#### 3.1.1 The PLN-LDA model

Informally, PLN-LDA assumes that (*i*) the sites belong to distinct and known classes, (*ii*) all sites in the same class have the same mean species abundances and (*iii*) those mean abundances may differ between classes but (*iv*) species interact in the same way in all classes. Formally, PLN-LDA for multivariate count data is a PLN model with an additional class covariate and a different decomposition of the mean vectors *µ*_*i*_. Assume the classes are labeled by *k* ∈ {1,…, *K*} and denote by *k*_*i*_ ∈ {1,…, *K*} the known class of site *i*. Finally, denote by 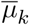 the mean vector in class *k*. The PLN-LDA model is the same as (1), where

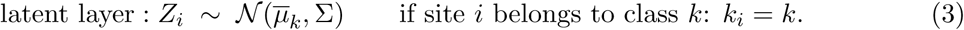

Compared to the standard PLN model (1), we need to estimate the additional class mean vectors 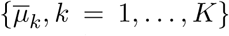. In presence of covariates, Model (3) can be extended to 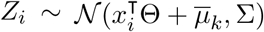.

##### Discriminant axes and prediction

Once estimated, the class means 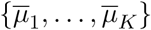 and reconstructed latent means *M* can be used to find axes of maximal discrimination of the classes in the latent layer, likewise in standard LDA. Scores along those axes can be used for visualization purposes and contributions of species to each axis can be used to identify systematic differences in abundances between the classes and potential biomarkers.

The classification of a new site from its species counts *Y*_new_ is based on Bayes’ rule. We first estimate the variational likelihood 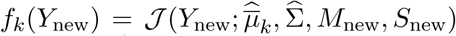 of observing counts *Y*_new_ if the new site was in class *k*. Note that 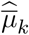 and 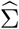 are extracted from the PLN-LDA fitted on the training sites whereas the variational parameters *M*_new_ and *S*_new_ must be optimized with respect to *Y*_new_. This corresponds to the VE step mentioned in Section 2. We then use an estimator 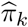 of the proportion of class *k* (typically the proportion of sites of class *k* among the learning sites). The *posterior* probability for the new site to belong to class *k* is then estimated as:

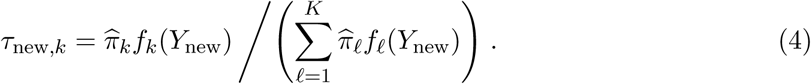

#### 3.1.2 Cow holobionts

To illustrate PLN-LDA, we consider the dataset introduced in Mariadassou *et al*. (2020) which consists of *n* = 256 bacterial communities sampled on three body sites (nose, mouth and vagina) of 45 primiparous PrimHolstein dairy cattle. The cattle comes from two divergent lineages, each sampled twice (1 month and 3 months after first delivery). Communities were sequenced using the hypervariable V3-V4 region of 16S rRNA as marker-gene. Sequences were cleaned and analyzed with DADA2 (Callahan *et al*., 2016) to create p = 1077 Amplicon Sequence Variants (ASVs). The aim is to assess whether the communities living in different body sites are different and how they differ.

We analyzed this data set by running PLN-LDA with time and lineage as covariates and body site as class. Offsets were computed as the log-total sums of counts over the 1077 ASVs, but we kept only 53 ubiquitous ASVs (with prevalence higher than 20% in at least one class) for the discriminant analysis. The 256 communities were split in two halves: a training test used to estimate the parameters and a test set used to assess the classification accuracy.

The results of our analysis with PLN-mixture are displayed in Figure 4: Panel *a*) shows that the first discriminant axis (LDA 1) separates vagina from nose and mouth whereas the LDA 2 separates nose from mouth. The inset correlation map shows the contribution of ASVs to LDAs: some ASVs are shared between nose and mouth or nose and vagina but almost none is shared between mouth and vagina. This is also obvious in the count matrix featured in panel *b*) where ASVs are reordered according to their position in the correlation map. The block structure indicates a strong association between some groups of species and body sites. Panel *c*) and *d*) show the same views for test samples. The inset confusion table of Panel *c*) shows a prediction accuracy of 95% (7 misclassified samples out of 128). The count matrices allow us to focus on the misclassified samples. 3 out of the 5 misclassified vagina samples have very small counts for all species. For those samples, the posterior probability of the second best class is around 0.25, indicating a quite high uncertainty. The misclassified nose (resp. mouth) sample is depleted in ubiquitous species typically found in other nose (resp. mouth) samples.

**Figure 4:**
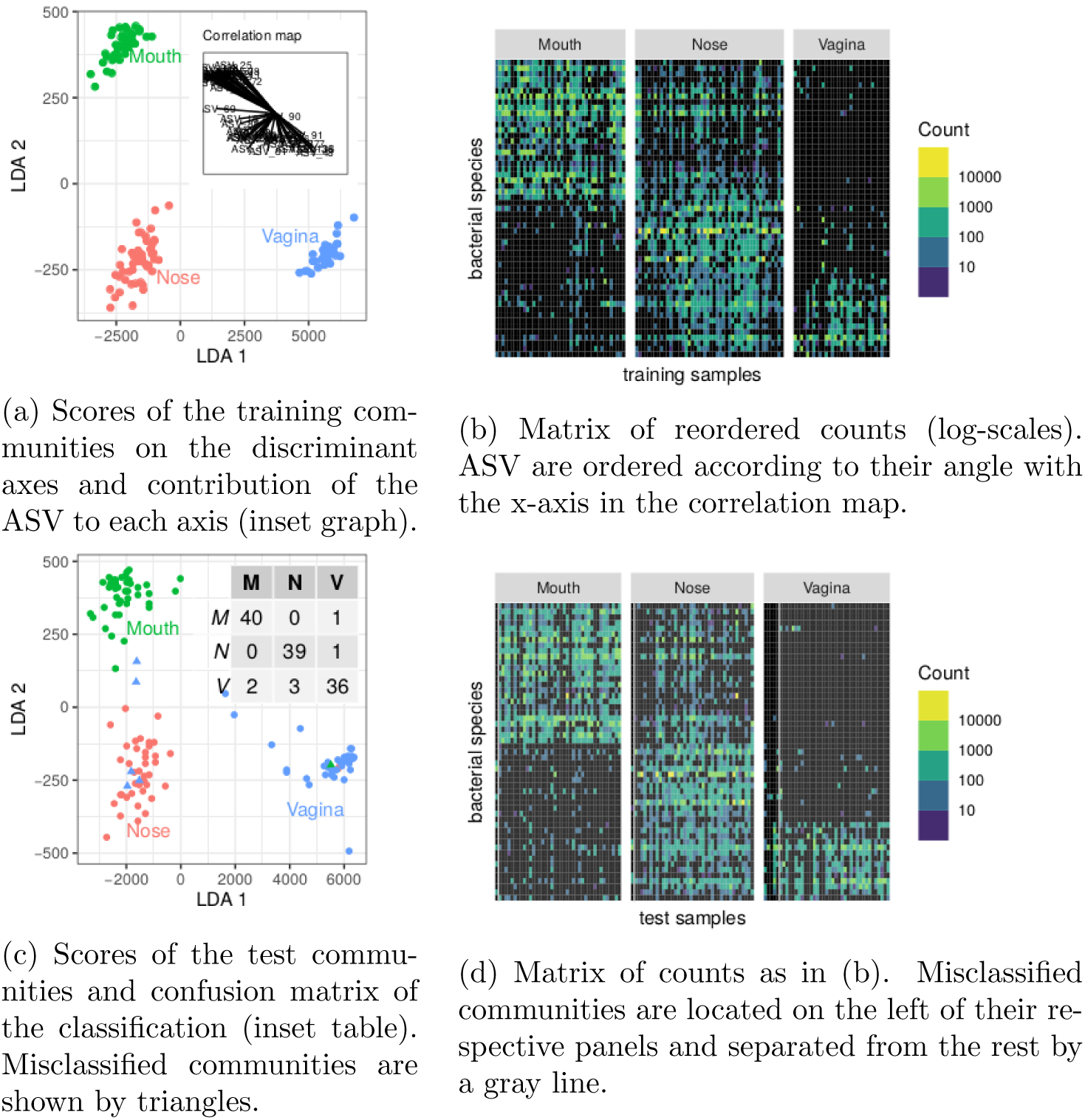
Result of the analysis of the cow holobionts data set with PLN-LDA: a few ASV are enough to find accurately discriminate the body sites.

### 3.2 Unsupervised classification with model-based clustering

This second variant is the analogue of Gaussian mixture models for Poisson-lognormal models. The objective is to perform model-based clustering on multivariate count tables, in order to find groups of homogeneous sites or samples in the data set.

#### 3.2.1 The PLN-mixture model

Informally, PLN-mixtures assumes that (*i*) the sites belong to *K unknown* groups, with different frequencies, (*ii*) all sites in the same group are homogenenous: they have the same mean species abundances and species interact in the same way in all sites, (*iii*) those mean abundances and interactions may differ between groups. Formally, PLN-mixture for multivariate count data is a PLN model with two latent layers: the first layer describes the (unknown) group membership of each site and the second layer embeds the distribution of the hidden site’s multivariate Gaussian vector conditional on its group membership. Note *K* the number of groups and *C*_*i*_ ∈ {1,…, *K*} the (unknown) group of site *i*. The PLN-mixture model assumes that each site has a probability *π*_*k*_ to belong to group *k*, so that *C*_*i*_ has a multinomial distribution. The latent vector *Z*_*i*_ associated with a site from group *k* is then assumed to have a multivariate Gaussian distribution with group-specific parameters 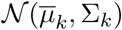. The latent layer of the original model (1) is therefore split as follows:

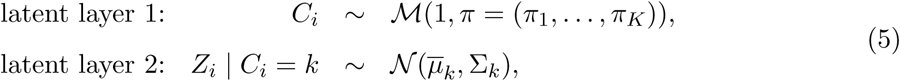

where the 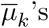 ’s and Σ_*k*_’s are the respective vector of means and the covariance matrix of the *K* components of the mixture, and π is the vector of the mixture proportions. Compared to the standard PLN model (1), we need to estimate the parameters {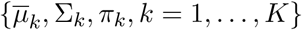, Σ_*k*_, *π*_*k*_, *k* = 1,…, *K*} as well as the group membership – or cluster – *C*_*i*_ of each sample. Covariates can also be included in the PLN-mixture model, changing the second layer of Eq. (5) into 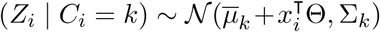. This extension is useful to correct for known environmental structuring factors and and recover some residual group structure among the sites.

The main difference between the PLN-LDA and PLN-mixture models is that the group (or class) memberships of the sites are known in the former, whereas they need to be inferred in the latter. The other difference is that the current implementation of PLN-mixture allows the covariance matrix Σ to vary across groups, whereas it is assumed to be constant in PLN-LDA. An important byproduct of the PLN-mixture model is the posterior probability, which can be used to actually classify sites into groups. These probabilities are iteratively computed along the algorithm using Eq. (4).

##### Parametrization of the covariance in PLN-mixture models

When using parametric mixture models like Gaussian mixture models, it is not recommended to consider general co-variance matrices Σ_*k*_ with no special restriction, especially when dealing with a large number of species. Indeed, the total number of parameters to estimate in the model can become prohibitive: in the general case, a PLN-mixture model with *K* components like in (5) has *K* × (*p*(*p* + 3)/2) model parameters, plus the *K* × 2(*n* × *p*) variational parameters. To reduce the computational burden and avoid over-fitting two different, more constrained parametrisations of the covariance matrices of each component are currently implemented in the `PLNmodels` package (on top of the general form of Σ_*k*_):

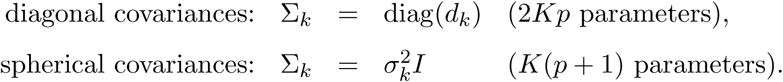

The diagonal structure assumes that, given the group membership of a site, all species abundances are independent. The spherical structure further assumes that all species have the same biological variability. In particular, in both parametrisations, all observed covariations are caused only by the group structure. For readers familiar with the `mclust R` package (Fraley & Raftery, 1999), which implements Gaussian mixture models with many variants of covariance matrices of each component, the spherical model corresponds to `VII` (spherical, unequal volume) and the diagonal model to `VVI` (diagonal, varying volume and shape). Using constrained forms of the covariance matrices enables PLN-mixture to provide a clustering even when the number of sites *n* remains of the same order, or smaller, than the number of species *p*.

#### 3.2.2 Oaks powdery mildew

To illustrate PLN-mixture, we consider the dataset introduced in Jakuschkin *et al*. (2016) which consists of microbial communities sampled on the surface of n = 116 oak leaves. Communities were sequenced with both the hypervariable V6 region of 16S rRNA as marker-gene for bacteria and the ITS1 as marker-gene for fungi. Sequences were cleaned, clustered at the 97% identity level to create OTUs and only the most abundant ones were kept (see Jakuschkin *et al*. (2016) for details) resulting in a total of *p* = 114 OTUs (66 bacterial ones and 44 fungal ones). One aim of this experiment is to understand the association between the abundance of the fungal pathogenic species *E. alphitoides*, responsible for the oak powdery mildew, and the other species. Furthermore, the leaves were collected on three trees with different resistance levels to the pathogen, which we call the tree *susceptibility*. We use this example to assess the ability of model-based clustering to recover, without feeding this information to the model, the existence of groups of leaves with different origins.

We analyzed this data set by running PLN-mixture for a number of component varying from 1 to 5. We selected the final number of components with a variant of the Integrated Classification Likelihood (ICL: Biernacki *et al*., 2000), tuned for our own PLN framework. Since the abundances were measured separately for fungi and bacteria, we define a different offset term *o*_*ij*_ for each OTU type to take into account the differences in sampling effort and marker genes. Offsets are still computed as the log-total sums of reads, including those of filtered out OTUs, for each OTU type.

The results of our analysis with PLN-mixture are displayed in Figure 5: Panel *a*) shows the evolution of the approximated log-likelihood (which is strictly increasing with the number of components, as expected) and the evolution of the ICL criterion, which suggest a model with 4 components. Panel *b*) displays a scatter-plot of the expected latent position 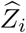, after performing simple PCA since pairs-to-pairs plot would be unreadable with *p* = 114 species. We also colored the site according to the most probable components according to our PLN-mixture, which shows that we recover the strong latent structure visible in the individual factor map. On Panel *c*), we compare the memberships of the series of PLN-mixture model with the tree susceptibility, by means of various measures for clustering comparison (ARI, AMI and NID, see Vinh *et al*., 2010). It then becomes obvious that the clustering found by PLN-mixture is highly related to the susceptibility level of each tree. Note that, even if apparently quite strong, this pattern in the data is not directly visible on the table of counts, as shown by the re-ordered version of the expected counts. This somewhat supports the modeling strategy of PLN-mixture and PLN in general, with a Poisson emission and a latent Gaussian layer.

**Figure 5:**
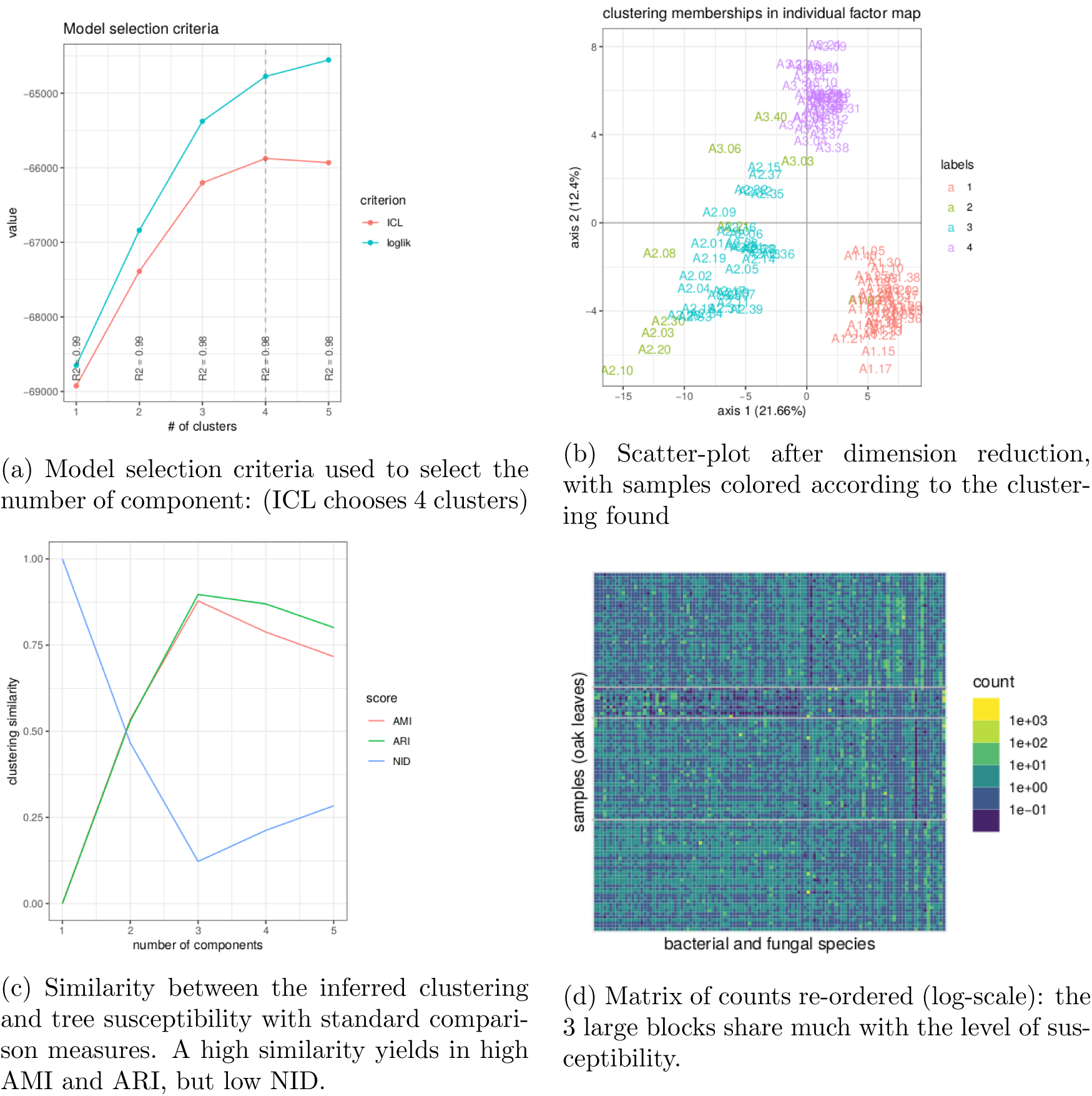
Result of the analysis of the oaks powdery mildew data set with PLN-mixture: the tree susceptibility to the pathogen is easily recovered, and seems to strongly structure the data set.

### 3.3 Dimension reduction with principal component analysis

We now turn to the extensions of the PLN model (1) that mostly deal with the modeling of the dependency between species, which is encoded in the covariance matrix Σ. One first way to depict species dependency is to look for a few underlying (that is: unknown) factors that may have have an impact on the whole community. The intuition behind this reasoning is that the p species actually respond to few unobserved drivers that structure most of their variations. As a consequence, finding such factors amounts at performing dimension reduction as it suggests that the variations of abundances can be summarized in a virtual space with much fewer dimensions than the number of species. This is especially desirable for studies involving a large number of species, when one looks for important pattern of diversity and tries to find structure in large data sets. This is exactly what principal component analysis (PCA) is designed for. We illustrate the PLN-PCA model by continuing the analysis started in the previous section on the oaks powdery mildew data set.

#### 3.3.1 Probabilistic Poisson PCA with PLN model

In the standard PLN model (1), the latent vector *Z* belongs to a latent space of dimension *p*, with one dimension per species. This assumes that any species can co-vary arbitrarily with any other, which allows for fine scale inferences but also becomes costly very quickly when the number of species *p* increases. PLN-PCA assumes instead the existence of *q* (with *q* » *p*) strong structuring unknown factors (*e.g.* environmnental filters) that govern the fluctuations of all species. All observed covariations between species then reflect those factors. Formally, PLN-PCA assumes that the latent vectors *Z*_*i*_ are fully determined by *q* structuring factors as follows:

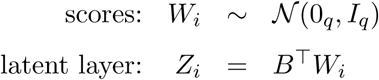

where the *W*_*i*_’s are supposed to be iid. As a consequence, the *Z*_*i*_’s are iid as well, with distribution

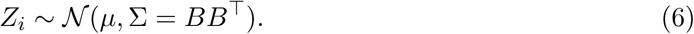

This is a strict extension of the probabilistic PCA model of Tipping & Bishop (1999) to the PLN model (Chiquet *et al*., 2018). The *p* × *q* matrix *B* is the analogue of the *rescaled* loadings in PCA: *B*_*jh*_ measures the impact of the *h*-th factor on the *j*-th species. Likewise, *W*_*i*_ is a *score* vector: *W*_*ih*_ is the value of the *h*-th factor for the *i*-th observation. The dimension *q* corresponds to the number of structuring factors, or equivalently to the number of axes in the PCA and the rank of Σ = *BB*^T^. The PLN-PCA model can thus be viewed as a PLN model with the low rank constraint rank(Σ) = *q* on the covariance matrix Σ. The number of parameters in the PLN-PCA model is (*p* + 1)*q*, down from *p*(*p* + 1)/2 in the standard model. Again, we can simply include covariates in the PLN-PCA model by changing Equation (6) into 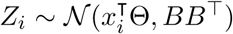. This is useful to correct for strong known structuring factors and to investigate weaker factors.

#### 3.3.2 Oaks powdery mildew

We continue the analysis of the oaks data set begun for PLN-mixture. As seen before, there is an obvious structure in the data explained by the tree susceptibility. Knowing this, one may be interested in exhibiting a remaining dependence structure that is not induced by the tree susceptibility.

In Panel *a*) of Figure 6, we represent the biplot for the first two principle components obtained with PLN-PCA, not accounting for the tree susceptibility, after selecting the best possible rank *q* thank to the ICL criterion. As expected after our clustering study with PLN-mixture, the factorial map exhibits a strong structure where individual are spread into three groups corresponding to the level of susceptibility of the trees where the leaves were sampled. We also projected the 10 species with the highest contribution to the first two principal components. Interestingly, the pathogen *E. alphitoides* point toward the group of susceptible trees, indicating that the presence of the disease is one of the main underlying drivers.

**Figure 6:**
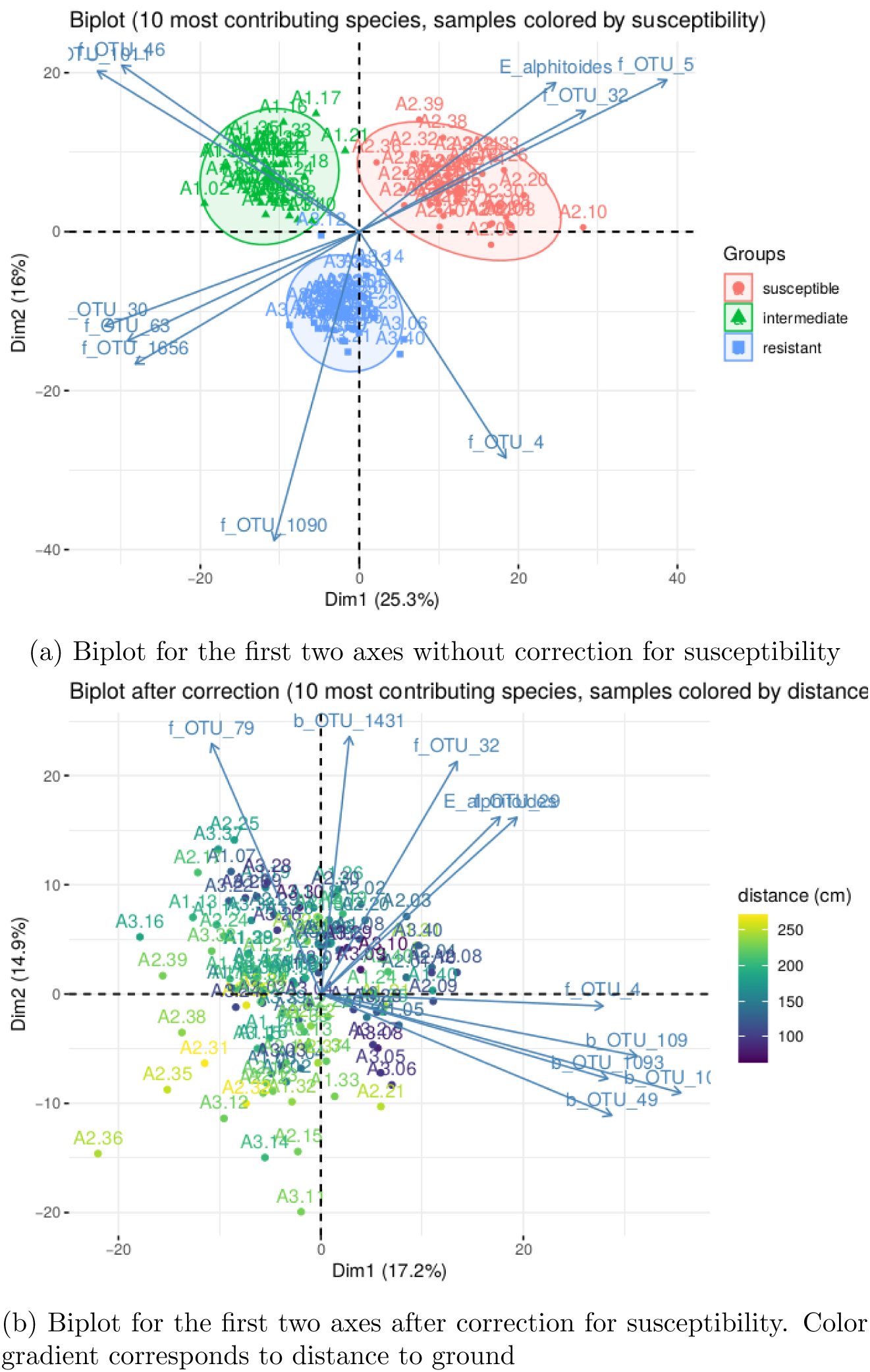
Result of PLN-PCA analysis on oaks powdery mildew data set

In Panel *b*) of Figure 6, we show how PLN-PCA can help exploring second-order structuring effects that are masked by strong first-order effect, that is here: the tree susceptibility. To do so, we included the susceptibility as a covariate to remove its effect and highlight a weaker effect: the map shows that the communities are structured by the distance of the leaf to the ground. The effect of covariates on the abundance of *E. alphitoides* were also consistent. When taking the susceptible tree as a reference, the estimated parameters *θ*_*ij*_ associated with the intermediate and resistant trees were respectively −3.94 (∼ 50 fold abundance decrease) and −7.05 (∼ 1000 fold abundance decrease).

### 3.4 Network inference

As a last extension of the PLN model, we introduce the analogue of graphical-Lasso (Banerjee *et al*., 2008; Friedman *et al*., 2008) for the inference of interaction networks. The ultimate goal is to find pairs of species that are in direct interaction. Direct interactions are hard to identify from covariation patterns in general as many different mechanisms can lead to the same patterns. For example, shared habitat preferences or reliance for growth on a metabolite produced by a third species, none of which requires interaction, (see e.g. Popovic *et al*., 2019), can lead to *statistical associations* that are indistinguishable from those of obligate symbiosis, an extreme form of direct interaction.

Formally, species can be associated but they are in direct interaction only if they are still *dependent* after conditioning on both the covariates (abiotic effects) and all the other species (biotic effects). In the Gaussian setting, this distinction coincides with the difference between *correlation* and *partial correlation*. Correlations between pairs of species is captured by the variance matrix Σ, whereas partial correlations are encoded by its inverse: the precision matrix Ω = [*ω*_*jk*_]_1≤*j,k*≤*p*_ = Σ^−1^. In this setting, species *j* and *k* are associated as soon as *σ*_*jk*_ ≠ 0 but are in direct interaction if and only if *ω*_*jk*_ ≠ 0 (Lauritzen, 1996).

#### 3.4.1 Network inference with the PLN model

The PLN network model for multivariate count data can be viewed as a PLN model with a constraint on the coefficients of Ω. Because the network is usually supposed to be sparse (*i.e.* only a few pairs of species are expected to be in direct interaction), we assume that the precision matrix Ω is sparse and the PLN-network model is the same as (1) with

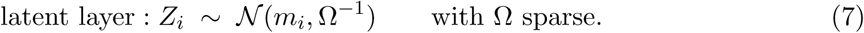

Both the PLN-PCA and the PLN-network models impose contraints on the covariance matrix Σ, but the low-rank constraint used in PLN-PCA aims at identifying few important unknown structuring factors, whereas the sparsity constraint used in PLN-Network aims at identifying direct interactions between species. Unlike previous extensions, this requires substantial modification of the objective function to be optimized, which becomes:

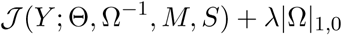

where |Ω|_1,0_ is the sum of the absolute values of the non-diagonal terms of Ω (diagonal terms are not penalized) and *λ* is a penalty coefficient. The term *λ*|Ω|_1,0_ forces many coefficients of 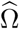 to be null. This is an extension of the graphical-Lasso (Banerjee *et al*., 2008; Friedman *et al*., 2008) to the PLN model (Chiquet *et al*., 2019). The parameter *λ* controls the number of edges in the network (larger *λ* yields fewer edges) and can be chosen in various ways, including model selection (Foygel & Drton, 2010) and resampling (Liu *et al*., 2010).

#### 3.4.2 Barents ftsh

We illustrate the use of the PLN-network model on the Barents fish dataset. We focus on the way the inferred network is modified when introducing covariates in the model. To this aim, we fitted the PLN-network model with (*a*) no covariates, (*b*) two environmental covariates (temperature and depth) and (*c*) all covariates (*i.e.* the previous two plus the geographical location) using a common *λ*-grid (with 20 values spaced equally in log scale between *λ*_min_ = 0.03 to *λ*_max_ = 15.17) for all models.

Figure 7 (top right panel) shows that the number of edges increases as the penalty decreases, as expected. It also shows that, for any penalty, the number of edges decreases as (plain lines) the richness of the number of edges increases (*c* > *b* > *a*) and that most edges recovered in the full (*c*) model are also recovered in the partial models (*a*, black dotted curve) and (*b*, blue dotted curve). This suggests that naive inference identifies not only genuine edges but also spurious ones corresponding to co-variations induced shared habitat preferences (captured here by temperature, depth and location). Interestingly, the dotted curve shows that the proportion of common edges between models *b* and *c* is higher than the one between models *a* and *c*. This suggests that environmental covariates rather than geographical location explain a substantial part of the apparent species co-variations.

**Figure 7:**
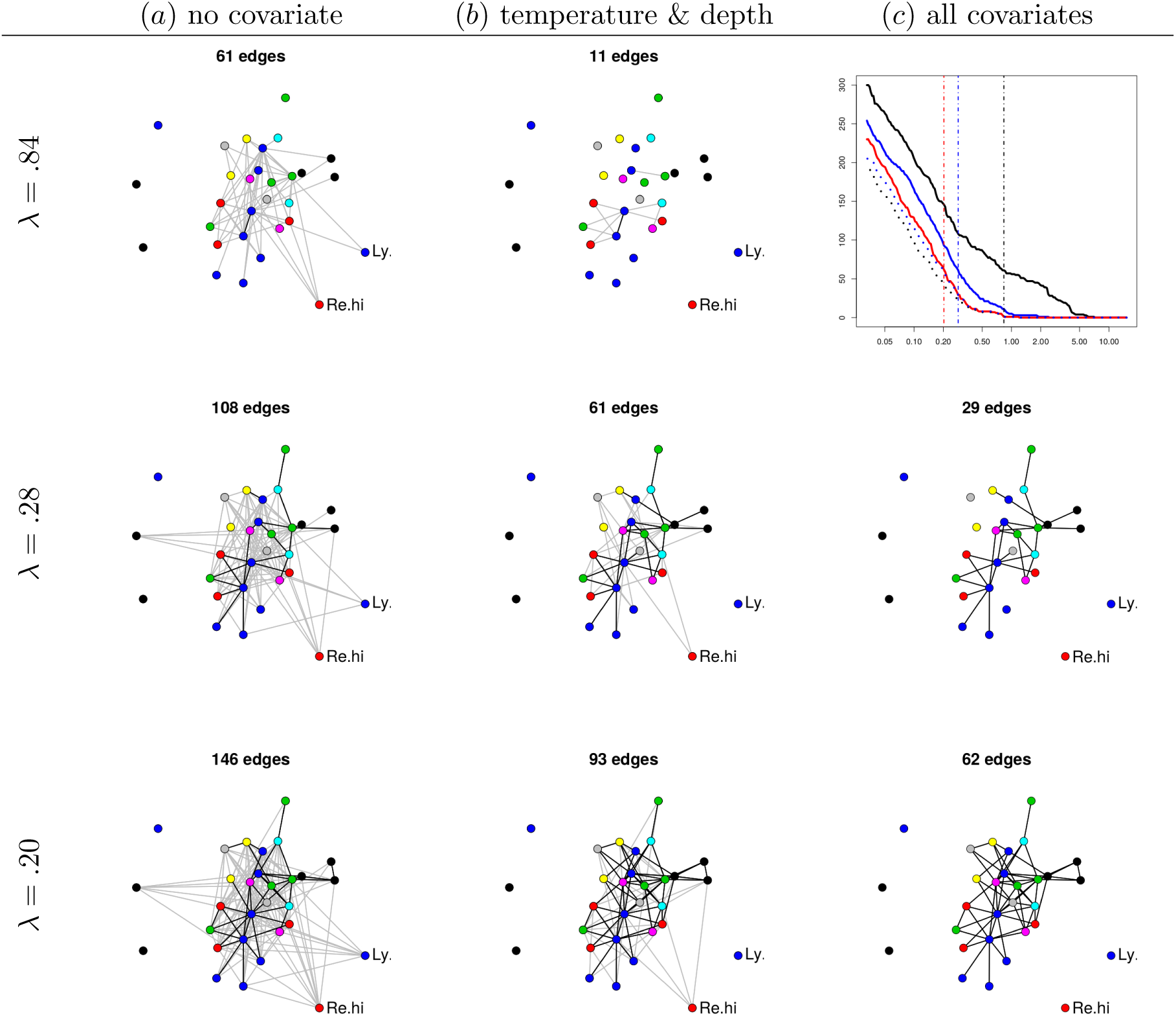
Inferred networks with increasing penalties (top: *λ* =. 84, middle: *λ* =. 28, bottom: *λ* =. 20) for different covariate sets (see top line). Each node corresponds to a given species. The position of the nodes are kept fixed. Node color: species family (see Fossheim *et al*., 2006). Black edges: edges in common with the network inferred with all covariates and same *λ*. The missing network (top right panel, all covariates and *λ* =. 84) contains only one edge. Top right: number of edges as a function of *λ*. Black: no covariate, blue: temperature and depth, red: all covariates, dotted black: common edges with no and all covariates, dotted blue: common edges between two and all covariates. Vertical dashed lines: the three chosen values of *λ*.

The rest of Figure 7 displays the networks inferred with the three models for three different levels of sparsity (controlled by *λ*). For an illustrative purpose, the values of *λ* have been chosen so that, in average, each species interacts with two others for each of the three models (*a*), (*b*) and (*c*). This results in networks with approximately 2*p* = 60 edges. One conclusion is that a set of core species seem to have direct interactions, or at least, interactions that cannot be simply explained by geographical location and environmental covariates (bottom right panel). On the contrary, some interactions seem to be actually indirect. For example, the interactions between the longear eelpout (*Ly.se*) and some species from the core group disappear when accounting for temperature and depth, suggesting that the covariation of their respective abundances results from shared environmental preferences. Similarly, the interactions between the Greenland halibut (*Re.hi*) and the core group is kept when correcting for temperature and depths, but disappears when correcting for location (longitude and latitude) suggesting that these interactions actually reflect a common response to fluctuations of biotic or abiotic characteristics across sites. To confirm this interpretation, we fitted an over-dispersed Poisson generalized linear model for the abundance of both species (not shown). We found that both temperature and depth have a significant effect on the abundance of the longear eelpout and that the longitude has a significant influence on the abundance of the Greenland halibut (all corresponding *p*-values being smaller than 10^−4^).

## 4 Discussion

### 4.1 PLNmodels package

All the variants of the PLN model presented in this paper are available as an `R/C++` package `PLNmodels`, distributed on the CRAN CRAN.R-project.org/package=PLNmodels. The package comes with a set of accompanying functions and methods for visualization and diagnostic. It also relies on the user-friendly GLM-like syntax to define all the models, so that users familiar with (generalized) linear models will feel at home. The development version is available on github.com/jchiquet/PLNmodels and all models are fully documented as vignettes available from the package website jchiquet.github.io/PLNmodels. The Barents and oaks mildew data sets analyzed in Sections 2 and 3 are included in the package.

### 4.2 Dedicated inference algorithms

We purposely avoided to enter into technical details, especially regarding the inference algorithms. Still, each extension of the PLN model illustrated above raises specific estimation issues. For all of them, a very naive solution would be to use the standard PLN model as a pre-processing step to retrieve the latent vectors *Z*_*i*_’s and then apply standard PCA, LDA, mixture and network inference to those vectors. Unfortunately, this solution is flawed: it does not propagate the uncertainty properly as the *Z*_*i*_ are estimated rather than known. The accuracy of the estimates comes precisely from the fact the the model parameters (Θ, Σ, *B*,…) are always estimated together with the latent or variational parameters (*M*, *S*, *τ*,…), which systematically leads to a complex high-dimensional optimization problem.

PLN-LDA and PLN-mixture can be recast as simple variants of the original PLN model and we can thus rely on the same inference algorithm, with a few minor modifications. In contrast, the inference algorithms of PLN-PCA and PLN-network models are quite more involved and detailed respectively in Chiquet *et al*. (2018) and Chiquet *et al*. (2019).

### 4.3 Future works

As shown along this paper, the Poisson-lognormal model provides a versatile framework for a large set of abundance data analyses. Thanks to its flexibility, many other extensions could be considered, either to include more sophisticated models or to account for data peculiarities. Two obvious examples come to mind. First, the basic PLN model assumes independence across sites, which mean that the spatial organization of the sites cannot be explicitly modeled, except through the recording of environmental descriptors as covariates. Adding spatial dependency would obviously be interesting, but requires methodological development as it would combine a sites’ dependence structure with the species’ dependence structure. The same obviously holds for times series of abundance data. Second, many experiments yield in a large proportion of null count, that cannot be explained by under-sampling alone. There is large literature (see Wagh & Kamalja, 2017, for a survey) devoted to distinguishing the *structural* zeroes (due to absent species) from the *sampling* zeroes (due to the combination of rare species and low sampling effort). A popular method, which can be adapted to PLN, is to consider Zero-Inflated distributions, where an additional latent layer codes for the presence/absence of each species at each site and absent species automatically lead to structural zeroes. As seen before, adding latent layers requires specific developments.

As shown in Section 2.2, the proposed estimation procedures yield accurate and unbiased estimates, but statistically grounded guarantees are still needed. Further theoretical analysis is required to get more insights both in terms of parameter and model uncertainty, especially confidence intervals. Several paths can be explored: (*i*) resampling procedures (which comes at a high computational cost), (*ii*) alternative estimation criterion like composite likelihoods (Varin *et al*., 2011) (for which statistical guaranties can be derived in a more systematic way) and (*iii*) the general theory of M-estimation (van der Vaart, 1998; Westling & McCormick, 2015), to which variational estimation belongs.

## Acknowledgments

This work was supported by the French ANR grants ANR-18-CE02-0010 Ecological Networks (EcoNet), ANR-17-CE32-0011 Next Generation Biomonitoring (NGB) and ANR-18-CE45-0023 Statistics and Machine Learning for Single Cell Genomics (SingleStatOmics). The authors thank Colin Fontaine for helpful comments and advices on early versions of the manuscript.

